# Blood draw site and blood matrix influence mineral assessment

**DOI:** 10.1101/2024.11.23.625018

**Authors:** Mikail G. Alejandro, Kathleen Schultz, David W. Killilea

**Affiliations:** Department of Pediatrics, University of California, San Francisco, CA, United States; Office of Research, University of California, San Francisco, CA, United States

**Author notes:** **Correspondence:** David W. Killilea.

**Keywords:** mineral assessment, sample processing, calcium, copper, iron, magnesium

## Abstract

Circulating mineral concentrations are used to assess nutritional status or as markers for clinical conditions, but the quality of these measurements depends on the methods used for blood processing. Since the procedures for measuring mineral levels in blood are not standardized, discrepancies in sampling may influence the analytical results. We previously reported that zinc content in blood samples showed significant variation depending on several pre-analytical variables selected during blood collection and processing. In this study, we extend our analysis to determine how other mineral levels might be affected by different blood draw sites (capillary or venous) or sample matrices (plasma or serum). Sequential capillary and venous blood samples were collected from a diverse cohort of sixty healthy adults and analyzed for multiple minerals using inductively coupled plasma– optical emission spectrometry. Calcium, copper, iron, and magnesium were the only minerals that were detected in all samples and were free from contamination in the blood collection tubes used for the study. When assessing different blood draw sites, the concentrations of calcium, copper, iron, and magnesium were 2-11% higher from capillary compared to the venous plasma. When assessing different blood sample matrices, the concentrations of calcium, copper, and magnesium were 2-5% higher in serum compared to plasma samples, whereas the concentration of iron was 7% higher in plasma compared to serum samples. The differences observed in these four essential minerals from discrepant draw sites and blood matrices demonstrate the importance of controlling key pre-analytic variables when assessing mineral levels in blood.

## 1. Introduction

Measuring mineral levels in the blood is a common clinical laboratory test, although only a few minerals (e.g. copper and zinc) are reliable indicators of their own nutritional status.^1,2^ This disconnect is known to result from homeostatic control and inflammation signals that can uncouple the circulating mineral levels from the actual nutritional state of the tissues. Yet the measurements of minerals within blood can still provide valuable information about other metabolic imbalances or specific pathological conditions. For example, blood calcium levels are held within a tight range even during dietary insufficiency, likely due the important roles that calcium plays in the vascular system. However, blood calcium levels are known to be abnormal in patients with hyperparathyroidism and chronic kidney disease, meaning the measurement of circulating calcium could have diagnostic utility.^3,4^ The methods for determining the mineral levels within the blood should be described by precise protocols, but unfortunately standardization is uncommon. Different approaches for mineral measurements may result in analytical discrepancies, prompting our interest to determine how circulating minerals respond to different pre-analytical variables during blood collection and processing.

One key pre-analytical variable is the draw site for blood samples. The most common method for blood draw is by phlebotomy from the venous circulation, typically at the cubital fossa.^5^ The blood is withdrawn from the vein using a syringe assembly that connects to specialized blood collection tubes. Phlebotomy has the advantage of being universally accepted, commonly used, and capable of yielding a large blood volume in a single session. However, phlebotomy does require some infrastructure, including trained technicians, sterile supplies, and a controlled clinical setting for safe collection of the blood sample; this increases the cost and complexity for a clinical study. The most common alternative to phlebotomy is a fingerstick using a single-use lancet with manual collection of blood droplets into specialized tubes.^6^ The fingerstick has the advantage of simplicity in requiring minimal training, low-cost supplies, and less risk of infection. However, the procedure yields substantially lower blood volume than a phlebotomy and draws blood from the capillary circulation, which is a mixture of arterial, venous, and interstitial fluid. A few older studies have reported differences in several metabolites between venous and capillary blood, but those reports were not widely appreciated, were conducted using older techniques, and often included few minerals. We recently reported that circulating levels of zinc were 8% higher in plasma from capillary compared to venous blood from the same donors.^7^ We were interested to see if other minerals would be affected by the blood draw site.

Another important variable for mineral assessment involves the type of blood matrix used for testing, specifically the use of plasma or serum fractions obtained from blood samples. Plasma contains all non-cellular constituents of the blood and can be separated immediately after the blood is drawn to reduce unwanted changes in mineral dynamics.^8^ However, plasma also requires the use of anticoagulant additives to block coagulation, which could potentially affect the distribution of the minerals within the blood. Alternatively, serum is similar to plasma but with the removal of clotting proteins, which has the advantage of not needing anticoagulants.^9^ However, serum samples require time for clotting during processing and collection tubes can include clotting activators as additives, both which might alter the distribution of the minerals within the blood. There is no clear consensus as to which matrix type is best for mineral assessment, so laboratories tend to choose the matrix that is more convenient for processing or follow historical preferences of the laboratory. Only a few studies have explored the differences in mineral concentrations between the plasma and serum, so additional attention to the impact of blood matrices on mineral assessment is needed.^9^ We recently reported that circulating levels of zinc were 5% higher in serum compared to plasma from the same donors.^7^ We were interested to see if other minerals would be affected by differences in the blood matrix.

The objective of this analysis is to determine how the concentrations of multiple minerals are influenced by blood draw site and blood matrix choices. The overall goal is to provide data to establish best practice guidelines for laboratories when processing blood samples for the purpose of mineral analysis.

## 2. Materials and Methods

### 2.1 Study Overview

This report is a secondary analysis of data collected in a previous study to assess the impact of specific pre-analytic variables on hemoglobin and zinc assessment in blood.^7,10^ The study was conducted completely on the campus of the Children’s Hospital Oakland Research Institute (CHORI), now part of the University of California San Francisco. The CHORI Institutional Review Board approved this study (2019-055). Sixty adults were recruited from Oakland, California, USA. Inclusion criteria included being 18 years or older, generally healthy with no chronic illness or blood disorders. Exclusion criteria included not meeting inclusion criteria or being pregnant due to known complexities in nutrient homeostasis. Participants were asked to avoid mineral and other supplements for 24 hours before blood draw and provide their age, gender, and race/ethnicity simply as markers of participant diversity. All demographic and anthropometric values were previously published.^10^ We utilized the Checklist for Reporting Stability Studies (CRESS) framework for comparability with other studies on the assessment of minerals from blood.^11^

### 2.2 Sample Processing

Participants provided non-fasting blood samples beginning with fingerstick(s) followed by conventional phlebotomy.^10^ Variables commonly associated with blood draw and processing were minimized, including use of the same clinical site, clinical procedures, phlebotomist, blood draw procedures, and sample handling procedures.^7,10^ Sequential blood collections were used to generate venous plasma, venous serum, and capillary plasma samples for each participant. The blood collection tubes (BCT) lot numbers, BCT additives, blood sample volumes, blood processing times, and detailed laboratory steps were previously published.^7,10^ The blood fractions were aliquoted into metal-free polypropylene tubes and stored at -70°C in a monitored ultralow freezer until ready for batch analysis.

### 2.3 Analysis of Mineral Content

Mineral content was determined by inductively coupled plasma optical emission spectrometry (ICP-OES). The ICP-OES methods and operating conditions were previously described and independently evaluated.^7,12^ In brief, 100 µl samples of plasma and serum were transferred to metal-free 15 ml conical tubes (Perfector Scientific), digested in 70% OmniTrace nitric acid (VWR) for 12-16 hours, and then diluted to 5% nitric acid with OmniTrace water (VWR). To test for mineral contamination, the BCTs and plasticware were rinsed with 5% nitric acid and transferred to conical tubes for analysis. All samples were vortexed 10-30 sec, centrifuged at 4,000 x*g* for 10 min, and then analyzed on a 5100 Synchronous Vertical Dual View ICP-OES (Agilent Technologies). The ICP-OES was calibrated using National Institute of Standards and Technology (NIST) traceable elemental standards and validated using Seronorm Trace Element Levels 1 and 2 standard reference materials (Sero AS) as external quality controls (**Supplemental Table 1**). Mean composite elemental values across 6 run days were within the reference range targets provided for Seronorm Trace Element Levels 1 and 2. Interassay precision of mineral content was assessed by testing 30 pooled plasma and 30 pooled serum samples generated from excess blood obtained from this study (**Supplemental Table 1**). For the pooled plasma, the CV was found to be 1.7% for calcium, 1.8% for copper, 2.2% for iron, and 1.8% for magnesium. For the pooled serum, the CV was found to be 3.8% for calcium, 2.7% for copper, 5.0% for iron, and 3.8% for magnesium.

### 2.4 Analysis of Hemolysis

Hemolysis was quantified by measuring the amount of hemoglobin released into the plasma or serum.^13^ Hemoglobin content was determined by a modified Drabkin’s assay according to previously published protocol.^14^ Each plasma or serum sample was measured in triplicate against calibration standards (0.5–360 mg/dL) from the same hemoglobin standard preparation, and previously reported.^10^

### 2.5 Data Analysis

Sample size was based on power calculations for differences in circulating zinc content as previously described.^7^ All graphing and statistical testing were conducted using Prism 10 (GraphPad Software, Inc). Outlier analysis was conducted using the GraphPad ROUT algorithm with Q=0.1%.^15^ Data normality was assessed with the D’Agostino-Pearson omnibus K2 normality test. For comparisons involving blood draw site or matrix, testing using Pearson correlation, Bland-Altman analysis, and two-tailed paired t test were used. For all analyses, statistical significance was assigned at p<0.05.

## 3. Results

### 3.1 Assessment of multiple minerals in blood samples

We previously used ICP-OES to determine how zinc measurements was influenced by several pre-analytical variables, but the instrument also records data for other minerals thus allowing for this secondary analysis.^7^ BCTs used for venous blood collection were selected based on their appropriateness for measuring zinc concentrations, but they were also certified for use with other trace elements including upper limits of mineral contamination for calcium (150 µg/L), copper (5 µg/L), iron (25 µg/L), and magnesium (40 µg/L), according to manufacturer’s data. To confirm this, we assessed several empty tubes from each lot of BCTs for background mineral content. Calcium, copper, iron, and magnesium were below detection in most of the empty tubes, whereas phosphorous, potassium, silicon, sodium, and sulfur were found to be moderately to heavily contaminated within the empty BCTs. Therefore, we limited our analysis to calcium, copper, iron, and magnesium within study blood samples. BCTs used for capillary blood collection had limited trace element certification, with limits for only lead at <1 ng/tube according to manufacturer’s data. Therefore, unused tubes from the same lot were tested by ICP-OES beforehand and were found to have undetectable levels of calcium, copper, iron, and magnesium.

### 3.2 Effect of blood draw site on mineral measurement

Calcium, copper, iron, and magnesium concentrations were quantified in blood drawn from capillary and venous sites from all study participants (**Table 1 and Fig 1**). All datasets passed normality tests, so parametric statistics were used.

**Table 1.**
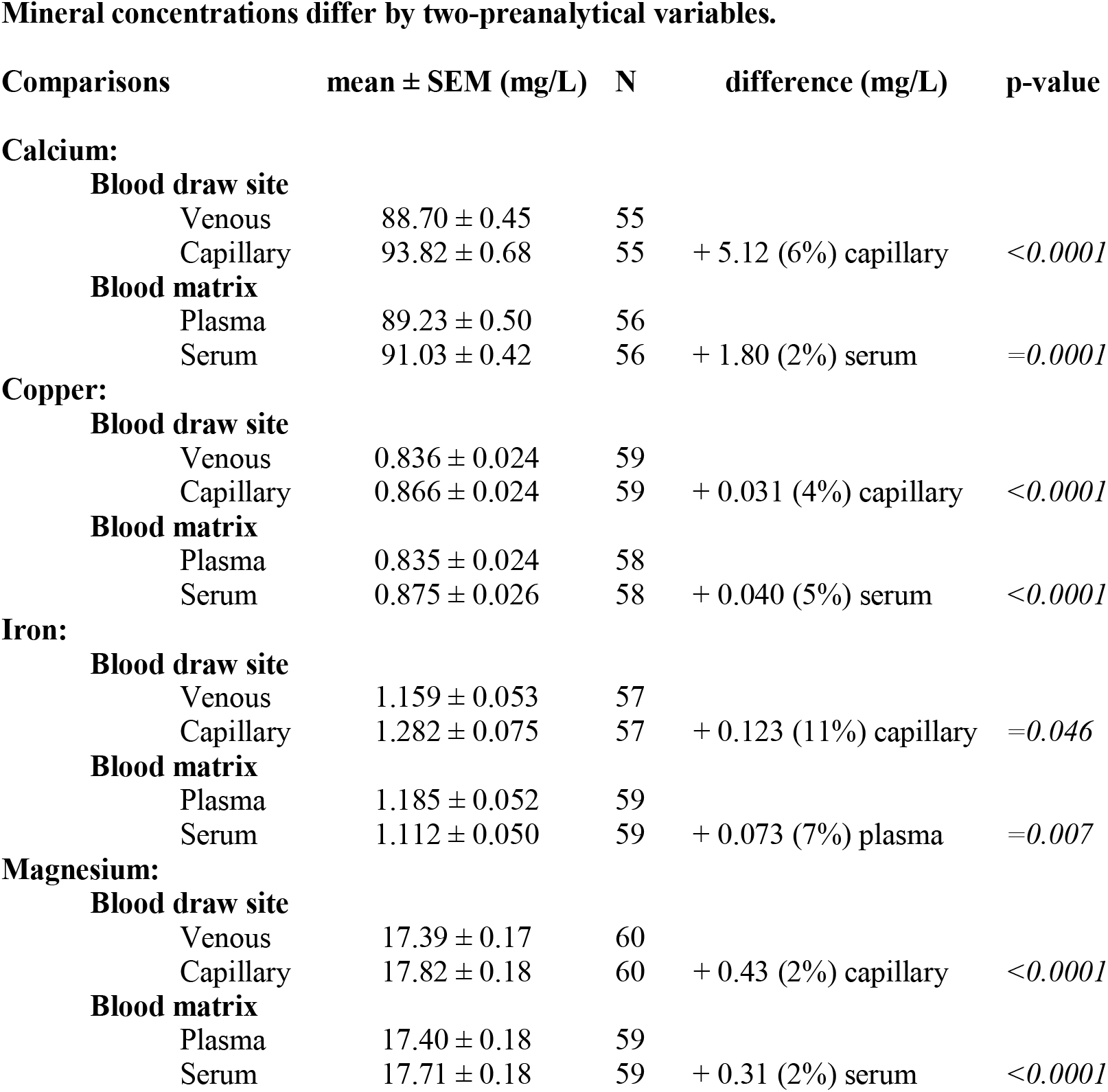
The mean ± SEM of the content for 4 minerals is shown as a function of specific pre-analytical variable. The number of samples measured (N), differences in measured mineral content, and p-values are listed.

**Mineral concentrations differ by two-preanalytical variables.**
| Comparisons | mean ± SEM (mg/L) | N | difference (mg/L) | p-value |
| --- | --- | --- | --- | --- |
| **Calcium:** |  |  |  |  |
| **Blood draw site** |  |  |  |  |
| Venous | 88.70 ± 0.45 | 55 | | |
| Capillary | 93.82 ± 0.68 | 55 | + 5.12 (6%) capillary | <0.0001 |
| **Blood matrix** |  |  |  |  |
| Plasma | 89.23 ± 0.50 | 56 | | |
| Serum | 91.03 ± 0.42 | 56 | + 1.80 (2%) serum | =0.0001 |
| **Copper:** |  |  |  |  |
| **Blood draw site** |  |  |  |  |
| Venous | 0.836 ± 0.024 | 59 | | |
| Capillary | 0.866 ± 0.024 | 59 | + 0.031 (4%) capillary | <0.0001 |
| **Blood matrix** |  |  |  |  |
| Plasma | 0.835 ± 0.024 | 58 | | |
| Serum | 0.875 ± 0.026 | 58 | + 0.040 (5%) serum | <0.0001 |
| **Iron:** |  |  |  |  |
| **Blood draw site** |  |  |  |  |
| Venous | 1.159 ± 0.053 | 57 | | |
| Capillary | 1.282 ± 0.075 | 57 | + 0.123 (11%) capillary | =0.046 |
| **Blood matrix** |  |  |  |  |
| Plasma | 1.185 ± 0.052 | 59 | | |
| Serum | 1.112 ± 0.050 | 59 | + 0.073 (7%) plasma | =0.007 |
| **Magnesium:** |  |  |  |  |
| **Blood draw site** |  |  |  |  |
| Venous | 17.39 ± 0.17 | 60 | | |
| Capillary | 17.82 ± 0.18 | 60 | + 0.43 (2%) capillary | <0.0001 |
| **Blood matrix** |  |  |  |  |
| Plasma | 17.40 ± 0.18 | 59 | | |
| Serum | 17.71 ± 0.18 | 59 | + 0.31 (2%) serum | <0.0001 |

**Figure 1:**
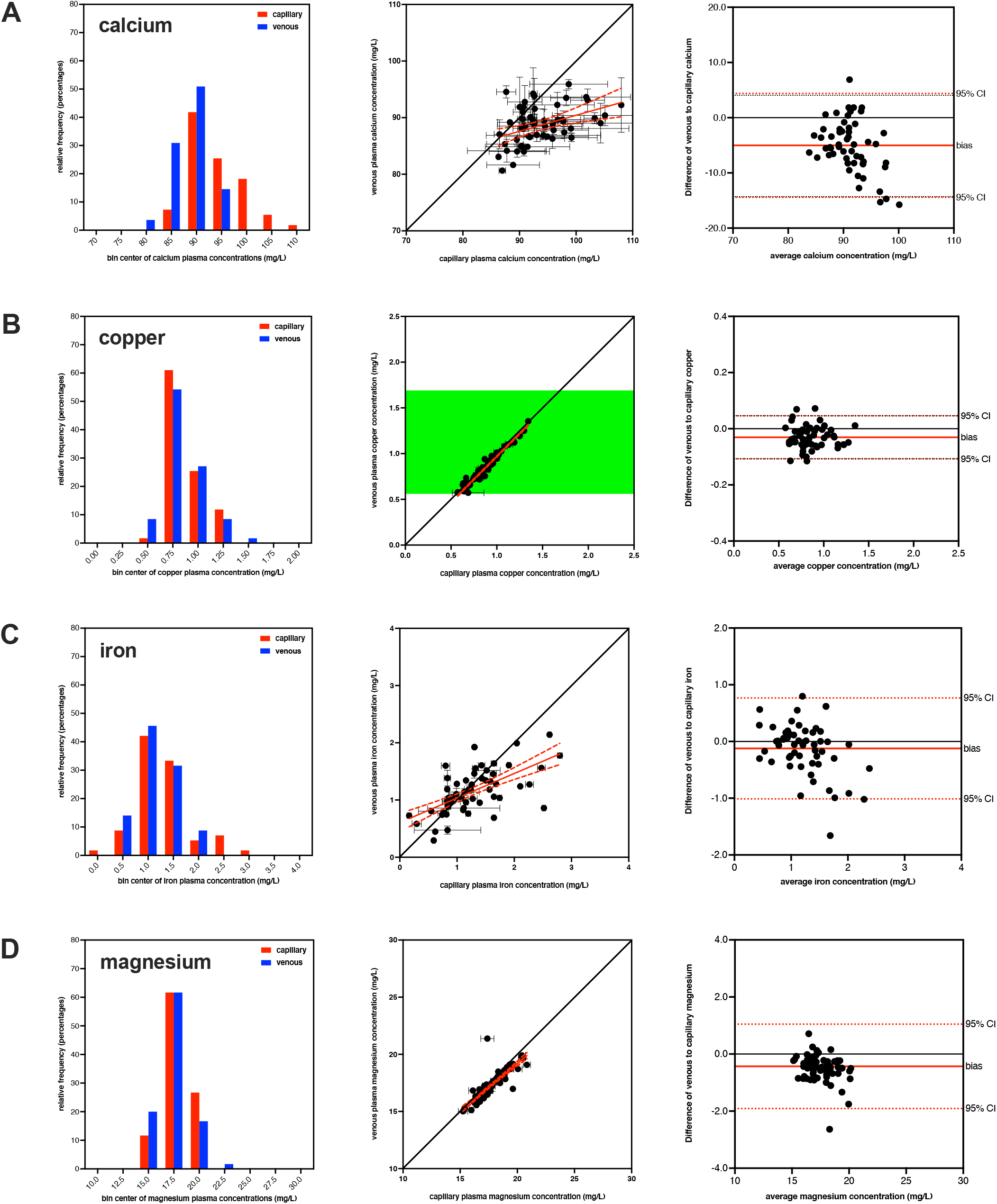
Comparison of mineral values from different blood draw site. The plasma concentrations of calcium (A), copper (B), iron (C), and magnesium (D), are shown from sequential blood draws taken from capillary and venous draw sites. The histograms in the first column show the distribution of mineral concentrations for capillary (red) and venous (blue) samples. The graphs in the second column show the correlation of the mineral concentrations (mean ± SEM) for both draw sites compared to the line of identity (solid black line) and clinical references ranges (green shading) where available.^16^ The linear correlation (solid red line) and 95% confidence band (dashed red line) of the mineral concentrations are also shown. Only copper had established reference ranges in venous plasma, while no reference ranges for minerals in capillary plasma were available. The graphs in the third column show the corresponding Bland-Altman assessment for both draw sites showing the line of identity (solid black line), mean bias (solid red line), and 95% confidence interval (dashed red line).

For calcium, the capillary samples had 3 sample identified as outliers while venous samples had 2 samples identified as outliers (with 0 overlap), resulting in a total of 55 matched samples. The mean calcium concentration was 93.82 ± 0.68 mg/L for capillary and 88.70 ± 0.45 mg/L for venous, with the difference being statistically significant (p<0.0001) using a two-tailed paired t-test. This indicated that capillary calcium concentration was elevated by 5.12 mg/L (6%) in comparison to venous blood values with a sequential blood draw from the same participants. Bland-Altman analysis revealed a similar bias of 5.13 mg/L for capillary compared to venous calcium concentrations.

For copper, the capillary samples had 1 sample identified as an outlier while venous samples had 1 sample identified as an outlier (with 1 overlap), resulting in a total of 59 matched samples. The mean copper concentration was 0.866 ± 0.024 mg/L for capillary and 0.836 ± 0.024 mg/L for venous, with the difference being statistically significant (p<0.0001) using a two-tailed paired t-test. This indicated that capillary copper concentration was elevated by 0.031 mg/L (4%) in comparison to venous blood values with a sequential blood draw from the same participants. Bland-Altman analysis revealed a similar bias of 0.031 mg/L for capillary compared to venous copper concentrations.

For iron, the capillary samples had 3 samples identified as outliers while venous samples had 0 samples identified as outliers, resulting in a total of 57 matched samples. The mean iron concentration was 1.282 ± 0.075 mg/L for capillary and 1.159 ± 0.053 mg/L for venous, with the difference being statistically significant (p=0.046) using a two-tailed paired t-test. This indicated that capillary iron concentration was elevated by 0.123 mg/L (11%) in comparison to venous blood values with a sequential blood draw from the same participants. Bland-Altman analysis revealed a similar bias of 0.123 mg/L for capillary compared to venous iron concentrations.

For magnesium, neither the capillary nor venous samples had outliers, resulting in a total of 60 matched samples. The mean magnesium concentration was 17.82 ± 0.18 mg/L for capillary and 17.39 ± 0.17 mg/L for venous, with the difference being statistically significant (p<0.0001) using a two-tailed paired t-test. This indicated that capillary magnesium concentration was elevated by 0.43 mg/L (2%) in comparison to venous blood values with a sequential blood draw from the same participants. Bland-Altman analysis revealed a similar bias of 0.43 mg/L for capillary compared to venous magnesium concentrations.

### 3.3 Effect of blood matrix on mineral measurement

Calcium, copper, iron, and magnesium concentrations were also determined within the same venous blood samples processed into plasma and serum from all study participants (**Table 1 and Fig 2**). All datasets passed normality tests, so parametric statistics were used.

**Figure 2:**
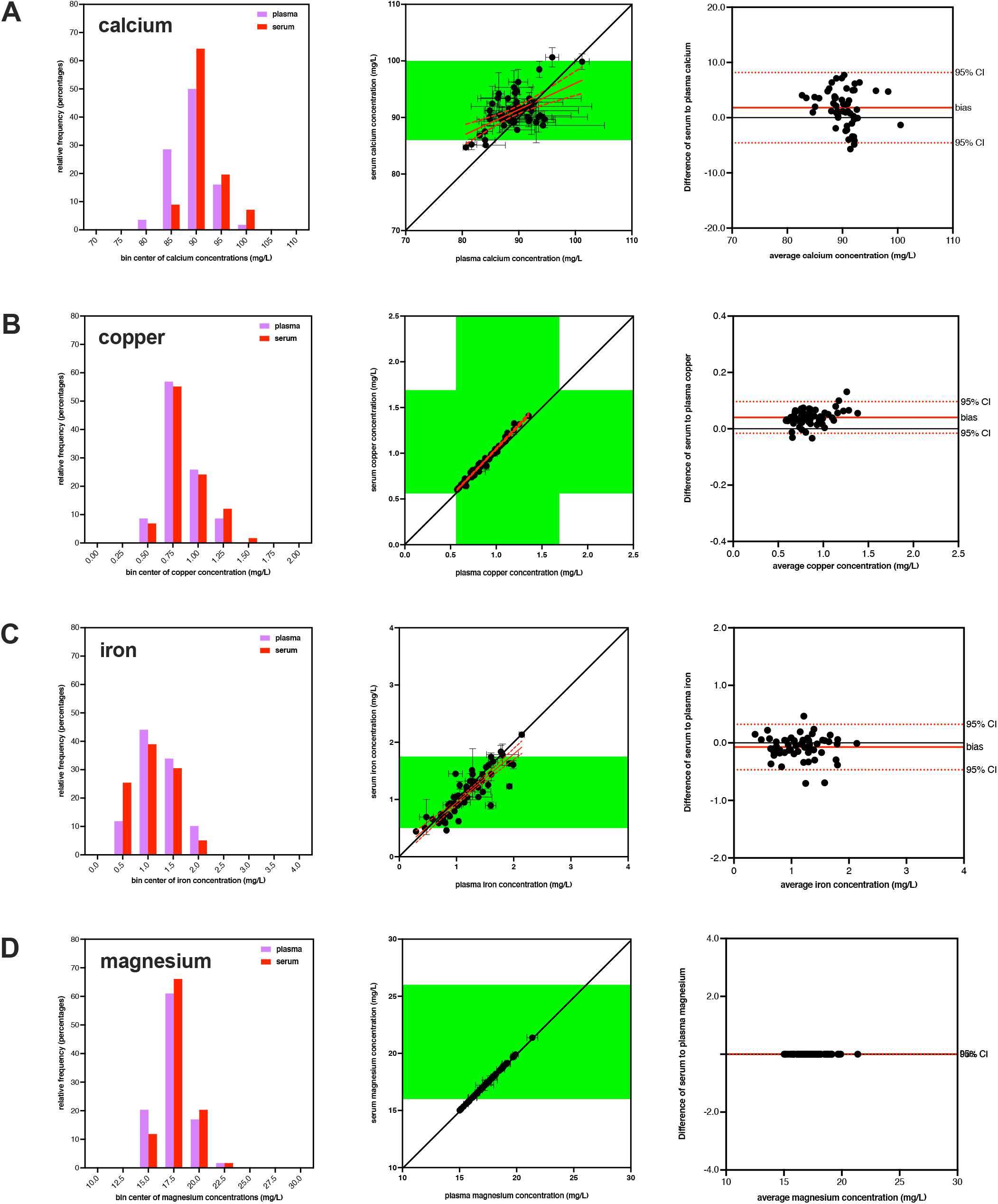
Comparison of mineral values from different blood matrices. The venous concentrations of calcium (A), copper (B), iron (C), and magnesium (D), are shown from the same blood draws but processed to plasma and serum matrices. The histograms in the first column shows the distribution of mineral concentrations for plasma (lavender) and serum (red) samples. The graphs in the second column show the correlation of the mineral concentrations (mean ± SEM) for both blood matrices compared to the line of identity (solid black line) and clinical references ranges (green shading) where available.^16^ The linear correlation (solid red line) and 95% confidence band (dashed red line) of the mineral concentrations are also shown. All found minerals had established reference ranges in venous serum, while only copper had established reference ranges in venous plasma. The graphs in the third column show the corresponding Bland-Altman assessment for both blood matrices showing the line of identity (solid black line), mean bias (solid red line), and 95% confidence interval (dashed red line).

For calcium, the plasma samples had 3 samples identified as outliers while serum samples had 3 samples identified as outliers and 1 missing value (with 3 overlap), resulting in a total of 56 matched samples. The mean calcium concentration was 89.23 ± 0.50 mg/L for plasma and 91.03 ± 0.42 mg/L for serum, with the difference being statistically significant (p=0.0001) using a two-tailed paired t-test. This indicated that serum calcium concentration was elevated by 1.80 mg/L (2%) in comparison to plasma values with a sequential blood draw from the same participants. Bland-Altman analysis revealed a similar bias of 1.80 mg/L for serum compared to plasma calcium concentrations.

For copper, the plasma samples had 1 sample identified as an outlier while serum samples had 1 sample identified as an outlier and 1 missing value (with 1 overlap), resulting in a total of 58 matched samples. The mean copper concentration was 0.835 ± 0.024 mg/L for plasma and 0.875 ± 0.026 mg/L for serum, with the difference being statistically significant (p<0.0001) using a two-tailed paired t-test. This indicated that serum copper concentration was elevated by 0.040 mg/L (5%) in comparison to plasma values with a sequential blood draw from the same participants. Bland-Altman analysis revealed a similar bias of 0.040 mg/L for serum compared to plasma copper concentrations.

For iron, the plasma samples had 1 sample identified as an outlier while serum samples had 0 samples identified as outliers and 1 missing value (with 1 overlap), resulting in a total of 59 matched samples. The mean iron concentration was 1.185 ± 0.052 mg/L for plasma and 1.112 ± 0.050 mg/L for serum, with the difference being statistically significant (p=0.007) using a two-tailed paired t-test. This indicated that plasma iron concentration was elevated by 0.073 mg/L (7%) in comparison to serum values with a sequential blood draw from the same participants. Bland-Altman analysis revealed a similar bias of 0.073 mg/L for plasma compared to serum iron concentrations.

For magnesium, the plasma samples had 1 sample identified as an outlier while serum samples had 0 samples identified as outliers and 1 missing value (with 1 overlap), resulting in a total of 59 matched samples. The mean magnesium concentration was 17.40 ± 0.18 mg/L for plasma and 17.71 ± 0.18 mg/L for serum, with the difference being statistically significant (p<0.0001) using a two-tailed paired t-test. This indicated that serum magnesium concentration was elevated by 0.31 mg/L (2%) in comparison to plasma values with a sequential blood draw from the same participants. Bland-Altman analysis had a bias of >0.001 mg/L for serum compared to plasma magnesium concentrations.

## 4 Discussion

We previously reported that the zinc levels in blood samples from healthy adults were influenced by several pre-analytical variables during blood collection and processing.^7^ In that study, zinc was measured using ICP-OES which has the capability to detect many minerals simultaneously, allowing for secondary analysis of other minerals from the previous dataset. Of the detectable minerals, four could be reliably quantified in all samples and were free of contamination in the BCTs, namely calcium, copper, iron, and magnesium. Other minerals could be measured including phosphorous, potassium, and sodium, but were found to have substantial background levels within the empty BCTs so they could not be included in this analysis. The four selected minerals were then tested to see how different pre-analytical variables would affect their measurement despite being from the same donor during the same blood draw.

Our results showed that calcium, copper, iron, and magnesium were all influenced by the choice of blood draw site and sample matrix. However, the direction and magnitude of responses mostly differed from the few similar studies previously reported in the literature for each pre-analytical variable. This was surprising at first, but there are a few possible reasons to explain these discrepancies. First and foremost, the type and quality of BCTs selected for mineral analysis work is critical. Most older studies did not use BCTs that were certified for trace mineral use or did not report in house testing to verify that the BCTs were free of minerals that could have compromised their measurements. This is a critical first step for assessments of this type, as we and others have shown that conventional BCTs can have substantial amounts of some minerals due to the anticoagulants or other additives.^17–20^ This is a particular concern for older studies because the reagents and supplies available at that time had less advanced manufacturing standards and used lower grades of chemicals.^20,21^ In our study, trace element certified BCTs were used for all venous plasma and serum blood samples with defined mineral backgrounds well below that found in blood. For capillary draws, we could not source trace element certified BCTs, so empty tubes from the same lot were carefully tested to verify that they were free of contamination for the minerals under investigation.

Differences in BCT types are another potential reason for discrepancies between our results and previous studies investigating different pre-analytical variables. For example, there are multiple anticoagulants that can be used in BCTs to isolate plasma. We chose BCTs with K2-EDTA, but other options include sodium heparin, lithium heparin, sodium citrate, or acid citrate dextrose. There are also multiple additives in BCTs to isolate serum. We chose additive-free tubes, but other tubes utilize coagulation promoter additives such as silicon microbeads. Additionally, isolating serum requires an incubation step during processing to allow the clotting process to finish, which is given as different times and temperatures depending on the BCT venders. The different additives and preparation protocols can all contribute to the discrepancies in mineral values between studies. Specific differences in tube types from previous reports compared to our study will be highlighted.

### 4.1 Calcium

Calcium is the most abundant mineral in the human body and best known as the key constituent in bones and teeth, but also is essential for muscle contraction, vascular function, neuronal activity, and signal transduction. Calcium adequacy is a public health concern in the USA as approximately 42% of the population remains below the estimated average requirement (EAR) for daily calcium intake.^2^ Normal levels for circulating calcium are between 82-102 mg/L for most adults, although those values are specified for serum and not plasma.^16^ Circulating calcium levels are under tight homeostatic control and do not typically reflect whole-body calcium status.^2^ Moreover, no clinical or biochemical test has been established which allows for direct determination of an individual’s calcium nutritional status, so the search for biochemical indicators of calcium status remains an active area of research. Measuring calcium levels in the blood may not be useful for determining nutritional status, but blood calcium levels are known to change under certain physiological or pathological states. Conditions that are associated with increased circulating calcium include hyperparathyroidism, Paget’s disease, and certain malignancies. ^3,4,22,23^ Conditions that are associated with decreased circulating calcium include chronic kidney disease, hypoparathyroidism, leprosy, hepatic cirrhosis, and vitamin D deficiency.^3,4,24,25^ Therefore, understanding the impact of pre-analytic variables on calcium assessment in blood has practical clinical value.

The impact of blood draw location on calcium levels has been previously evaluated. Early studies tested the concentrations of numerous blood constituents between capillary and venous samples, finding no significant differences in calcium levels.^26,27^ Later groups reported that calcium levels were slightly higher in venous compared to capillary serum sample, but judged the differences to not be clinically relevant.^28,29^ In contrast, we found that calcium was significantly higher in capillary compared to venous plasma. In the Falch study, few details of the BCTs types were provided beyond than the vender, and no tube certification or background testing was provided. And in the Kupke *et al* study, no details of the types of BCTs or background testing were provided.

The impact of blood sample matrix on calcium levels was previously explored in only a few papers. Three studies have reported that calcium measurements were modestly elevated in plasma compared to serum.^30–32^ However, we found that calcium was significantly higher in serum compared to plasma. In the Herbert study, few details of the BCTs types were provided other than sodium citrate was stated as the anticoagulant for plasma samples. In the Lester and Varghese study, lithium heparin was stated as the anticoagulant for the plasma samples, but no information was given about the serum tubes. In the Linder *et al* study, EDTA was stated as the anticoagulant for plasma samples so likely the same as the K2-EDTA we used in our study, but there was no information given for the serum tubes. And none of these older studies reported certification or testing of tubes for background calcium levels.

### 4.2 Copper

Copper is a trace mineral with essential roles in energy production, connective tissue maintenance, oxidative stress defense, signal transduction, and neuronal function. The amount of copper needed for good health is modest and typically adequate in the average Western diet.^33^ Normal levels for circulating copper are given as between 0.7-1.4 mg/L for plasma and 0.56-1.69 mg/L for serum in most adults.^16^ Copper levels in the blood are influenced by changes in inflammation and other minerals, but circulating copper is still considered reflective of whole body copper status.^34^ Beyond the use as a nutritional marker, copper levels in the blood are also altered during specific clinical conditions. Conditions that are associated with increased circulating copper include rheumatoid arthritis, anemia, Hodgkin’s lymphoma, and pregnancy.^35–38^ Conditions that are associated with decreased circulating copper include Wilson’s disease, Menkes’ syndrome, and Alzheimer’s disease.^39–42^ Therefore, it is important to know the impact of pre-analytic variables on copper assessment.

The impact of blood draw location on copper levels was explored in at least one study. Rodríguez-Saldaña *et al* tested several nutrient and non-nutrient metal levels in blood taken from different blood sites, but observed no significant differences in copper content.^43^ In contrast, our findings showed that copper was significantly higher in capillary compared to venous blood. Rodríguez-Saldaña *et al* did report the use of trace metal certified K2-EDTA tubes for venous samples, but there was no description of certification or background testing of copper levels in their capillary tubes.

The impact of blood sample matrix on copper levels was also previously reported by two teams, though neither directly tested for differences between the plasma versus serum values of copper.^44,45^ As a result, it is unclear if there were any statistical differences in copper levels between samples matrices in these studies. Additionally, neither study reported use of certified tubes or tested for background copper, though the Toro-Raman study did state the BCTs were ‘metal free.” Our results showed that copper was significantly higher in serum compared to plasma.

### 4.3 Iron

Iron is the most abundant trace metal in the body and is essential for oxygen transport, energy production, immune function, oxidative stress defense, and neuronal function. Iron adequacy is a public health concern in the USA, particularly for women of reproductive age.^33^ While only 1% of the population is listed as below the EAR for daily iron intake, the true prevalence of iron-deficiency anemia in the USA may be closer to 14-19%.^46,47^ Normal levels for iron in the blood are given as between 0.5-1.75 mg/L for adults, although those values are specifically listed for serum and not plasma.^16^ Most of the iron in the blood is bound to the protein transferrin, so the iron saturation of transferrin is a more robust indicator of iron status than blood iron content alone. Furthermore, the circulating levels of iron are under homeostatic control and easily overridden by inflammatory signals, as the sequestration of circulating iron is a central protective mechanism during infection.^33^ Measuring total iron levels in the blood may not be optimal for determining nutritional status, but iron levels in the blood are known to change under certain physiological or pathological states.

Conditions that are associated with increased circulating iron include acute kidney disease, acute leukemia, and several hemoglobinopathies.^48–50^ Conditions that are associated with decreased circulating iron include chronic kidney disease, hypothyroidism, and infection.^51–53^ Therefore, understanding the impact of pre-analytic variables on iron assessment in blood could hold practical clinical value.

The impact of blood draw location on iron levels was also explored in one study. Rodríguez-Saldaña *et al* tested several nutrient and non-nutrient metal levels in blood taken from different blood sites, but observed no significant iron in copper content. ^43^ In contrast, our findings show that iron measurements were significantly elevated in capillary blood compared to venous blood. Rodríguez-Saldaña *et al* did report the use of trace metal certified K2-EDTA tubes for venous samples, but there was no description of certification or background testing for iron levels in their capillary tubes.

The impact of blood sample matrix on iron levels was also previously reported. Two teams reported that iron levels were higher in serum than plasma.^44,54^ However, our findings showed that iron measurements were significantly elevated in plasma over serum. In the Kasperek *et al* study, heparin was stated as the anticoagulant for plasma samples, but no information on serum tubes. The study did report the background iron levels in the plasma tubes, but not the serum tubes. This study also stated that the iron levels being higher in serum were ‘well established’ but we would dispute that claim. In the Jablan *et al* study, sodium citrate was used as the anticoagulant in plasma tubes and silica microparticles was used as an additive for serum tubes. Neither study reported certification or background testing for iron levels in their capillary tubes.

### 4.4 Magnesium

Magnesium is an essential mineral that cofactor in hundreds of enzymes involved in energy production, bone health, nucleic acid synthesis, protein synthesis, ion transport, oxidative stress defense, and signal transduction. Magnesium adequacy is a public health concern in the USA as approximately 51% of the population remains below the EAR for daily magnesium intake.^2^ Normal levels for magnesium in the blood are given as between 16-26 mg/L for adults, although those values are specifically listed for serum and not plasma.^16^ Circulating magnesium levels are under tight homeostatic control and do not typically reflect the whole-body magnesium status.^2^ Moreover, no clinical or biochemical test has been established which allows for direct determination of an individual’s magnesium nutritional status; as with calcium, the search for biochemical indicators of magnesium status remains an active area of research. Measuring magnesium levels in the blood may not be useful for determining nutritional status, but blood magnesium levels are known to change under certain physiological or pathological states. Conditions that are associated with increased circulating magnesium include hypothyroidism, Addison’s disease, and chronic kidney disease.^3,55,56^ Conditions that are associated with decreased circulating magnesium include hyperthyroidism, acute pancreatitis, chronic alcoholism, and sometimes pregnancy.^57–61^ Therefore, understanding the impact of pre-analytic variables on magnesium assessment in blood could hold practical clinical value.

The impact of blood draw location on magnesium levels was also explored in at least one previous study. Matthiesen and colleagues reported a greater ionized magnesium concentration in venous blood compared to capillary blood, but this is distinct from total magnesium levels and thus not comparable to our analysis.^62^ Our findings were that magnesium levels were elevated in capillary blood compared to venous.

The impact of blood sample matrix on magnesium levels was also explored in at least one previous paper. Linder and colleagues reported that magnesium values were higher in plasma than in serum.^31^ However, our findings were that magnesium levels were elevated in serum compared to plasma. In the Linder *et al* study, EDTA was stated as the anticoagulant for plasma samples so likely same as K2-EDTA as used in our study, but there was no information on serum tubes. And none of these older studies reported certification or background testing of tubes for magnesium levels.

## 5. Conclusion

We measured the concentrations of four essential minerals as a function of two different pre-analytical variables to identify whether process discrepancies could influence the observed circulating values. We found that both the draw site (capillary or venous) and sample matrix (plasma or serum) significantly affected the measurements of calcium, copper, iron, and magnesium in blood samples from healthy adults. These findings add to our previous report showing that circulating zinc levels were affected by draw site and sample matrix, illustrating the importance of controlling pre-analytical variables within a study. Additionally, these findings often differ with previous reports in the literature. However, those older reports tend to use different BCTs and/or fail to demonstrate the BCTs were clear of background contamination, resulting in concern with that reported differences in mineral levels.

Strengths of this study include the strict control over the parameters to remove extraneous influences that could confound the assessment of the specific pre-analytical variables of interest. Additionally, we collected blood samples from a diverse adult population to maximize applicability to communities within the USA. Limitations of this study include that the highly controlled conditions we utilized do not reflect the complexities found within a field assessment or large cohort study. Additionally, the statistical significance found for the mineral levels assessed in this study do not necessarily imply clinical significance as well. Some of the differences in mineral content were small, so it is not clear if they would affect the actual interpretation of mineral levels in a population. To both points, we believe that understanding the variability contingent within these pre-analytic variables is a necessary first step to select the best methodology for future studies utilizing mineral measurements.

## Data availability statement

The raw data supporting the conclusions of this article will be made available by the authors, without undue reservation.

## Ethics statement

The studies involving humans were approved by the Children’s Hospital Oakland Research Institute Institutional Review Board. The studies were conducted in accordance with the local legislation and institutional requirements. Written informed consent for participation in this study was provided by all participants.

## Author contributions

MGA: Data curation, Formal analysis, Writing—review & editing. KS: Data curation, Formal analysis, Investigation, Methodology, Project administration. DWK: Conceptualization, Data curation, Formal analysis, Investigation, Methodology, Project administration, Resources, Supervision, Funding acquisition, Validation, Writing—original draft.

## Acknowledgements

The authors wish to thank Nyla Sepulveda for phlebotomy assistance and thank Bonny Alvarenga, Tatiana Cheong, Darryl Chow, and Wesley Kwong for technical assistance.

