## Supplemental Table 1 for "Blood draw site and blood matrix influence mineral assessment"

**(1) ICP METHOD DETAILS**

| element | Ca | Cu | Fe | K | Mg | Na | P | S |
|---|---|---|---|---|---|---|---|---|
| wavelength (nm) | 373.69 | 324.754 | 238.204 | 766.491 | 280.27 | 588.995 | 213.618 | 181.972 |
| RF power (kW) | 1.2 | 1.2 | 1.2 | 1.2 | 1.2 | 1.2 | 1.2 | 1.2 |
| plasma flow rate (L/min) | 12 | 12 | 12 | 12 | 12 | 12 | 12 | 12 |
| nebulizer flow rate (L/min) | 0.7 | 0.7 | 0.7 | 0.7 | 0.7 | 0.7 | 0.7 | 0.7 |
| auxiliary flow rate (L/min) | 1 | 1 | 1 | 1 | 1 | 1 | 1 | 1 |
| minimum detection (mg/L) | 0.005 | 0.005 | 0.005 | 0.05 | 0.05 | 0.05 | 0.05 | 0.05 |
| maximum detection (mg/L) | 50 | 5 | 5 | 50 | 50 | 50 | 50 | 50 |

**(2) POOLED PLASMA DATA**

| element | Ca | Cu | Fe | K | Mg | Na | P | S |
|---|---|---|---|---|---|---|---|---|
| sample 1 | 101.095 | 1.123 | 1.996 | 1243.488 | 20.176 | 3102.648 | 133.066 | 6864.924 |
| sample 2 | 103.748 | 1.164 | 1.931 | 1282.903 | 21.003 | 3185.959 | 133.268 | 7050.922 |
| sample 3 | 106.440 | 1.173 | 1.940 | 1314.842 | 21.349 | 3266.383 | 136.234 | 7266.500 |
| sample 4 | 105.177 | 1.171 | 1.910 | 1305.533 | 20.998 | 3244.481 | 136.363 | 7239.050 |
| sample 5 | 105.875 | 1.173 | 1.930 | 1308.389 | 20.662 | 3254.619 | 133.529 | 7305.229 |
| sample 6 | 105.257 | 1.165 | 1.914 | 1307.777 | 21.075 | 3245.910 | 133.725 | 7300.505 |
| sample 7 | 105.439 | 1.169 | 1.925 | 1310.803 | 21.086 | 3248.806 | 132.848 | 7387.894 |
| sample 8 | 104.764 | 1.182 | 1.930 | 1301.810 | 21.049 | 3231.686 | 137.415 | 7305.305 |
| sample 9 | 104.720 | 1.175 | 1.904 | 1304.927 | 20.837 | 3236.756 | 132.629 | 7339.937 |
| sample 10 | 104.867 | 1.165 | 1.900 | 1298.624 | 20.826 | 3226.930 | 132.782 | 7392.001 |
| sample 11 | 99.435 | 1.114 | 1.789 | 1234.913 | 19.661 | 3062.788 | 126.425 | 7076.540 |
| sample 12 | 104.041 | 1.164 | 1.894 | 1294.376 | 20.751 | 3212.544 | 128.364 | 7361.405 |
| sample 13 | 104.895 | 1.174 | 1.927 | 1306.045 | 20.982 | 3240.151 | 137.186 | 7487.938 |
| sample 14 | 102.975 | 1.144 | 1.869 | 1279.253 | 20.466 | 3177.831 | 132.247 | 7362.154 |
| sample 15 | 106.079 | 1.179 | 1.934 | 1314.646 | 21.064 | 3261.925 | 133.603 | 7595.071 |
| sample 16 | 103.717 | 1.162 | 1.876 | 1286.849 | 20.593 | 3197.359 | 129.226 | 7526.599 |
| sample 17 | 104.173 | 1.165 | 1.892 | 1296.085 | 21.020 | 3214.179 | 124.902 | 7566.395 |
| sample 18 | 104.895 | 1.176 | 1.914 | 1303.894 | 20.715 | 3231.458 | 125.303 | 7649.468 |
| sample 19 | 104.785 | 1.181 | 1.902 | 1306.217 | 20.756 | 3235.608 | 130.354 | 7636.232 |
| sample 20 | 105.007 | 1.166 | 1.912 | 1307.527 | 20.595 | 3243.677 | 134.001 | 7684.627 |
| sample 21 | 103.401 | 1.155 | 1.877 | 1289.548 | 20.720 | 3194.696 | 131.460 | 7531.464 |
| sample 22 | 104.770 | 1.159 | 1.900 | 1308.946 | 20.834 | 3242.830 | 133.910 | 7718.311 |
| sample 23 | 105.273 | 1.170 | 1.893 | 1302.947 | 20.852 | 3225.829 | 129.403 | 7715.306 |
| sample 24 | 105.416 | 1.175 | 1.940 | 1307.108 | 20.855 | 3234.653 | 128.130 | 7761.443 |
| sample 25 | 99.832 | 1.103 | 1.838 | 1239.612 | 20.005 | 3077.047 | 135.829 | 7349.448 |
| sample 26 | 104.074 | 1.157 | 1.873 | 1294.332 | 20.696 | 3246.703 | 136.290 | 7720.964 |
| sample 27 | 105.623 | 1.173 | 1.905 | 1311.104 | 20.926 | 3240.998 | 132.435 | 7824.726 |
| sample 28 | 104.431 | 1.170 | 1.891 | 1302.966 | 20.741 | 3223.890 | 131.121 | 7755.604 |
| sample 29 | 102.881 | 1.154 | 1.859 | 1283.152 | 20.583 | 3180.013 | 126.396 | 7683.446 |
| sample 30 | 100.078 | 1.118 | 1.800 | 1245.887 | 19.912 | 3087.330 | 125.140 | 7446.203 |
| mean | 104.100 | 1.161 | 1.899 | 1293.000 | 20.730 | 3209.000 | 131.800 | 7464.000 |
| standard deviation | 1.815 | 0.020 | 0.041 | 22.840 | 0.372 | 55.630 | 3.697 | 235.000 |
| standard error of mean | 0.331 | 0.004 | 0.008 | 4.170 | 0.068 | 10.160 | 0.675 | 42.910 |
| coefficient of variation | 1.74% | 1.76% | 2.17% | 1.77% | 1.80% | 1.73% | 2.81% | 3.15% |

**(3) POOLED SERUM DATA**

| element | Ca | Cu | Fe | K | Mg | Na | P | S |
|---|---|---|---|---|---|---|---|---|
| sample 1 | 95.420 | 1.172 | 1.320 | 191.897 | 18.908 | 3132.038 | 124.199 | 7455.555 |
| sample 2 | 100.281 | 1.223 | 1.353 | 201.612 | 20.144 | 3280.877 | 142.572 | 7897.348 |
| sample 3 | 100.569 | 1.228 | 1.381 | 203.962 | 20.357 | 3312.084 | 135.430 | 8003.470 |
| sample 4 | 100.916 | 1.239 | 1.393 | 202.570 | 20.165 | 3288.474 | 136.089 | 7972.826 |
| sample 5 | 99.722 | 1.215 | 1.357 | 201.447 | 20.225 | 3280.377 | 136.341 | 7937.153 |
| sample 6 | 95.377 | 1.180 | 1.314 | 192.086 | 19.135 | 3182.486 | 132.890 | 7582.914 |
| sample 7 | 100.812 | 1.231 | 1.364 | 203.254 | 20.121 | 3298.987 | 135.929 | 7995.864 |
| sample 8 | 100.789 | 1.240 | 1.384 | 202.874 | 20.416 | 3352.454 | 133.853 | 8063.261 |
| sample 9 | 100.430 | 1.226 | 1.366 | 203.097 | 20.150 | 3307.082 | 134.443 | 8043.489 |
| sample 10 | 92.961 | 1.147 | 1.267 | 187.079 | 18.527 | 3059.490 | 128.521 | 7464.656 |

| | | | | | | | | |
|---|---|---|---|---|---|---|---|---|
| sample 11 | 95.222 | 1.166 | 1.346 | 189.819 | 19.018 | 3148.655 | 130.896 | 7539.609 |
| sample 12 | 98.179 | 1.204 | 1.343 | 198.828 | 19.887 | 3279.075 | 135.167 | 7920.672 |
| sample 13 | 99.566 | 1.224 | 1.362 | 202.390 | 20.196 | 3323.076 | 136.402 | 7996.398 |
| sample 14 | 100.141 | 1.231 | 1.371 | 202.421 | 20.052 | 3333.629 | 135.808 | 8056.707 |
| sample 15 | 100.188 | 1.234 | 1.374 | 202.668 | 20.381 | 3297.650 | 131.428 | 8100.873 |
| sample 16 | 95.573 | 1.177 | 1.317 | 191.561 | 19.255 | 3135.640 | 130.072 | 7748.788 |
| sample 17 | 98.862 | 1.214 | 1.363 | 199.660 | 19.841 | 3286.969 | 133.575 | 8039.283 |
| sample 18 | 99.268 | 1.210 | 1.374 | 200.481 | 19.948 | 3250.170 | 135.448 | 8078.354 |
| sample 19 | 100.824 | 1.231 | 1.375 | 203.783 | 20.315 | 3297.985 | 135.274 | 8230.930 |
| sample 20 | 98.663 | 1.218 | 1.348 | 199.550 | 19.714 | 3232.234 | 132.761 | 8070.138 |
| sample 21 | 104.007 | 1.241 | 1.649 | 199.562 | 20.586 | 3293.540 | 140.066 | 8485.173 |
| sample 22 | 104.590 | 1.243 | 1.423 | 200.288 | 20.788 | 3301.359 | 135.298 | 8520.455 |
| sample 23 | 103.503 | 1.239 | 1.462 | 198.230 | 20.738 | 3266.080 | 136.319 | 8466.284 |
| sample 24 | 105.107 | 1.245 | 1.439 | 200.400 | 21.288 | 3314.030 | 141.285 | 8593.650 |
| sample 25 | 101.521 | 1.217 | 1.381 | 192.487 | 20.300 | 3187.993 | 140.472 | 8270.917 |
| sample 26 | 105.577 | 1.259 | 1.430 | 201.749 | 20.880 | 3298.480 | 146.768 | 8585.820 |
| sample 27 | 110.000 | 1.298 | 1.498 | 210.343 | 21.956 | 3436.013 | 145.446 | 8927.301 |
| sample 28 | 106.989 | 1.272 | 1.459 | 204.008 | 21.500 | 3358.733 | 139.412 | 8741.500 |
| sample 29 | 106.312 | 1.271 | 1.442 | 203.793 | 21.217 | 3341.731 | 140.508 | 8704.710 |
| sample 30 | 101.753 | 1.211 | 1.398 | 191.980 | 20.308 | 3167.878 | 145.208 | 8318.627 |
| mean | 100.800 | 1.224 | 1.388 | 199.500 | 20.210 | 3268.000 | 136.300 | 8127.000 |
| standard deviation | 3.841 | 0.032 | 0.069 | 5.307 | 0.762 | 80.450 | 5.054 | 378.500 |
| standard error of mean | 0.701 | 0.006 | 0.013 | 0.969 | 0.139 | 14.690 | 0.923 | 69.110 |
| coefficient of variation | 3.81% | 2.65% | 5.01% | 2.66% | 3.77% | 2.46% | 3.71% | 4.66% |

**(4) SERONORM DATA - INDIVIDUAL DATA**

| element | **Ca** | **Cu** | **Fe** | **K** | **Mg** | **Na** | **P** | **S** |
|---|---|---|---|---|---|---|---|---|
| Seronorm Level 1 | | | | | | | | |
| Day 1 | | | | | | | | |
| replicate 1 | 91.947 | 0.986 | 1.256 | 122.793 | 16.555 | 2695.524 | 35.360 | 986.899 |
| replicate 2 | 88.131 | 0.973 | 1.200 | 120.860 | 16.137 | 2660.288 | 34.807 | 997.842 |
| replicate 3 | 85.442 | 0.940 | 1.137 | 116.644 | 15.408 | 2569.471 | 33.400 | 980.328 |
| Day 2 | | | | | | | | |
| replicate 1 | 86.476 | 0.920 | 1.275 | 117.649 | 16.341 | 2592.278 | 39.290 | 950.993 |
| replicate 2 | 85.488 | 0.923 | 1.208 | 116.645 | 16.082 | 2539.339 | 39.255 | 962.982 |
| replicate 3 | 87.817 | 0.951 | 1.225 | 122.547 | 16.665 | 2691.941 | 41.030 | 1007.324 |
| Day 3 | | | | | | | | |
| replicate 1 | 96.017 | 0.942 | 1.507 | 118.117 | 16.182 | 2626.994 | 42.655 | 1202.940 |
| replicate 2 | 88.392 | 0.932 | 1.310 | 116.532 | 15.859 | 2581.947 | 33.538 | 1185.512 |
| replicate 3 | 90.617 | 0.927 | 1.282 | 115.929 | 15.871 | 2575.554 | 33.370 | 1243.050 |
| Day 4 | | | | | | | | |
| replicate 1 | 83.457 | 0.920 | 1.292 | 118.253 | 15.704 | 2532.525 | 33.040 | *NA* |
| replicate 2 | 82.567 | 0.923 | 1.271 | 119.059 | 15.819 | 2513.623 | 33.221 | *NA* |
| replicate 3 | 84.825 | 0.953 | 1.286 | 122.245 | 16.185 | 2581.958 | 34.092 | *NA* |
| Day 5 | | | | | | | | |
| replicate 1 | 83.261 | 0.927 | 1.316 | 116.297 | 15.718 | 2585.779 | 32.780 | 827.553 |
| replicate 2 | 85.757 | 0.930 | 1.320 | 117.875 | 15.764 | 2605.724 | 33.355 | 846.420 |
| replicate 3 | 90.602 | 0.983 | 3.066 | 123.304 | 16.371 | 2710.005 | 34.629 | 865.787 |
| Day 6 | | | | | | | | |
| replicate 1 | 85.511 | 0.946 | 1.286 | 114.907 | 15.850 | 2591.979 | 34.379 | 926.128 |
| replicate 2 | 87.892 | 0.964 | 1.275 | 115.538 | 16.196 | 2619.297 | 35.058 | 944.176 |
| replicate 3 | 86.106 | 0.950 | 1.245 | 115.728 | 16.093 | 2598.219 | 35.090 | 954.928 |
| Seronorm Level 2 | | | | | | | | |
| Day 1 | | | | | | | | |
| replicate 1 | 128.373 | 1.697 | 1.745 | 215.057 | 32.200 | 3211.186 | 94.097 | 1456.652 |
| replicate 2 | 136.869 | 1.711 | 1.736 | 217.655 | 32.690 | 3235.525 | 94.882 | 1476.821 |
| replicate 3 | 124.018 | 1.688 | 1.701 | 215.139 | 32.389 | 3199.745 | 94.364 | 1493.714 |
| Day 2 | | | | | | | | |
| replicate 1 | 119.471 | 1.620 | 1.731 | 207.722 | 32.096 | 3076.609 | 95.548 | 1340.842 |
| replicate 2 | 141.915 | 1.668 | 1.807 | 215.473 | 33.041 | 3152.867 | 99.022 | 1381.643 |

| | | | | | | | | |
|---|---|---|---|---|---|---|---|---|
| replicate 3 | 165.976 | 1.691 | 1.833 | 217.554 | 33.992 | 3177.909 | 99.836 | 1401.909 |
| Day 3 | | | | | | | | |
| replicate 1 | 134.717 | 1.614 | 1.896 | 206.190 | 32.329 | 3090.437 | 99.570 | 1845.173 |
| replicate 2 | 118.316 | 1.606 | 1.825 | 203.367 | 31.481 | 3049.710 | 89.969 | 1815.908 |
| replicate 3 | 118.129 | 1.608 | 1.844 | 203.529 | 31.529 | 3046.288 | 89.941 | 1868.804 |
| Day 4 | | | | | | | | |
| replicate 1 | 117.359 | 1.648 | 1.845 | 216.949 | 32.388 | 3057.366 | 95.886 | *NA* |
| replicate 2 | 120.412 | 1.719 | 1.869 | 223.366 | 33.072 | 3173.576 | 93.986 | *NA* |
| replicate 3 | 120.171 | 1.685 | 1.855 | 223.565 | 33.234 | 3160.290 | 94.223 | *NA* |
| Day 5 | | | | | | | | |
| replicate 1 | 126.388 | 1.706 | 1.972 | 220.643 | 33.040 | 3212.548 | 93.968 | 1244.522 |
| replicate 2 | 137.987 | 1.632 | 1.865 | 209.980 | 31.393 | 3089.736 | 89.587 | 1191.543 |
| replicate 3 | 123.800 | 1.702 | 1.913 | 221.431 | 32.973 | 3219.597 | 93.960 | 1255.337 |
| Day 6 | | | | | | | | |
| replicate 1 | 121.076 | 1.679 | 1.799 | 208.187 | 32.656 | 3161.267 | 93.501 | 1356.788 |
| replicate 2 | 121.920 | 1.661 | 1.775 | 206.960 | 32.416 | 3136.586 | 93.447 | 1386.108 |
| replicate 3 | 120.713 | 1.662 | 1.783 | 208.693 | 32.791 | 3153.189 | 93.476 | 1400.712 |

**(5) SERONORM DATA - SUMMARIZED DATA**

| element | **Ca** | **Cu** | **Fe** | **K** | **Mg** | **Na** | **P** | **S** |
|---|---|---|---|---|---|---|---|---|
| Seronorm Level 1 | | | | | | | | |
| Day 1 | 88.507 | 0.966 | 1.198 | 120.099 | 16.033 | 2641.761 | 34.522 | 988.356 |
| Day 2 | 86.594 | 0.931 | 1.236 | 118.947 | 16.363 | 2607.852 | 39.858 | 973.766 |
| Day 3 | 91.675 | 0.934 | 1.366 | 116.859 | 15.971 | 2594.832 | 36.521 | 1210.500 |
| Day 4 | 83.616 | 0.932 | 1.283 | 119.852 | 15.903 | 2542.702 | 33.451 | *NA* |
| Day 5 | 86.540 | 0.947 | 1.917 | 119.159 | 15.951 | 2633.836 | 33.588 | 846.587 |
| Day 6 | 86.503 | 0.953 | 1.269 | 115.391 | 16.047 | 2603.165 | 34.843 | 941.744 |
| mean | 87.239 | 0.944 | 1.378 | 118.385 | 16.044 | 2604.025 | 35.464 | 992.191 |
| standard deviation | 2.678 | 0.014 | 0.270 | 1.860 | 0.165 | 35.137 | 2.420 | 133.939 |
| Seronorm Level 1 reference range | | | | | | | | |
| minimum | 69 | 0.852 | 1.17 | 101 | 13.4 | 2330 | 43.3 | *NA* |
| target | 86 | 1.066 | 1.47 | 127 | 16.8 | 2917 | 54.2 | 981 |
| maximum | 104 | 1.281 | 1.77 | 153 | 20.1 | 3504 | 65.1 | *NA* |
| Seronorm Level 2 | | | | | | | | |
| Day 1 | 129.753 | 1.699 | 1.727 | 215.950 | 32.426 | 3215.485 | 94.447 | 1475.729 |
| Day 2 | 142.454 | 1.660 | 1.790 | 213.583 | 33.043 | 3135.795 | 98.136 | 1374.798 |
| Day 3 | 123.721 | 1.610 | 1.855 | 204.362 | 31.780 | 3062.145 | 93.160 | 1843.295 |
| Day 4 | 119.314 | 1.684 | 1.856 | 221.294 | 32.898 | 3130.410 | 94.698 | *NA* |
| Day 5 | 129.392 | 1.680 | 1.917 | 217.351 | 32.468 | 3173.960 | 92.505 | 1230.467 |
| Day 6 | 121.236 | 1.667 | 1.785 | 207.947 | 32.621 | 3150.347 | 93.475 | 1381.202 |
| mean | 127.645 | 1.666 | 1.822 | 213.414 | 32.539 | 3144.691 | 94.404 | 1461.098 |
| standard deviation | 8.399 | 0.031 | 0.067 | 6.259 | 0.444 | 50.985 | 2.001 | 230.928 |
| Seronorm Level 2 reference range | | | | | | | | |
| minimum | 95 | 1.538 | 1.72 | 176 | 27.1 | 2820 | 88 | *NA* |
| target | 119 | 1.925 | 2.15 | 221 | 33.9 | 3531 | 110 | 1335 |
| maximum | 143 | 2.312 | 2.58 | 265 | 40.7 | 4241 | 132 | *NA* |
